# Development of *Candida auris* microsatellite typing and its application on a global collection of isolates

**DOI:** 10.1101/792663

**Authors:** Theun de Groot, Ynze Puts, Indira Berrio, Anuradha Chowdhary, Jacques F. Meis

## Abstract

*Candida auris* is a pathogenic yeast that causes invasive infections with high mortality. Infections most often occur in intensive care units of healthcare facilities. It is crucial to trace the source and prevent further spread of *C. auris* during an outbreak setting, therefore, genotyping of *C. auris* is required. To enable fast and cost-effective genotyping, we developed a microsatellite typing assay for *C. auris*.

Short tandem repeats (STRs) in *C. auris* were identified, and a novel STR typing assay for *C. auris* was developed using 4 panels of three multiplex PCRs. Having shown that the microsatellite typing assay was highly reproducible and specific, a robust set of 444 *C. auris* isolates was investigated to identify genotypic diversity. In concordance with whole-genome sequencing (WGS) analysis we identified five major different *C. auris* clusters, namely, South-America, South-Asia, Africa, East-Asia and Iran. Overall, a total of 40 distinct genotypes were identified, with the largest variety in the East Asian clade. Comparison with WGS demonstrated that isolates with <20 SNPs are mostly not differentiated by STR analysis, while isolates with 30 or more SNPs usually have differences in one or more STR markers.

Altogether, a highly reproducible and specific microsatellite typing assay for *C. auris* was developed, which distinguishes the five different *C. auris* clades in identical fashion to WGS, while most isolates differing >20 SNPs, as determined via WGS, are also separated. This new *C. auris* specific genotyping technique is a rapid, reliable, cost-effective alternative to WGS analysis to speedily investigate outbreaks.

**Importance:** *Candida auris* is an emerging fungal pathogen now recognized as a threat to public health. The pathogen has spread worldwide and mainly causes hospital associated outbreaks. To track and trace outbreaks and to relate them to new introductions from elsewhere, whole genome sequencing and amplified fragment length polymorphism (AFLP) have been used for molecular typing. While the former is costly and only available in few centers, AFLP is a complicated technique and standardization is not possible. We describe a novel simple microsatellite genotyping technique based on small tandem repeats in the *C. auris* genome. Further we show that this microsatellite based genotyping technique has been proven comparable to WGS. Overall, this work provides a novel, rapid, reliable and cost-effective method of molecular outbreaks investigations of *C. auris*.

## Introduction

*Candida auris* is a pathogenic yeast which was first isolated in Korea in 1996 and first reported in 2009 in a Japanese patient who had a *C. auris* infection of the external ear canal.(1,2) In the following decade this yeast was found in locations all over the world, including South-Africa, India, Pakistan, Kuwait, Venezuela, UK and USA(3-8), suggesting its emergence might be a consequence of climate change.(9) Fungemia and wound infections are the most common clinical entities due to *C. auris*, which could subsequently lead to infection of other organs like heart and brain.(10) The mortality rate in patients with *C. auris* infections, which often already experience severe disease, reaches levels of 30-60%.(10, 11) A serious complication in the treatment of patients is the resistance of *C. auris* for multiple antifungal agents, such as fluconazole and voriconazole.(12) Most *C. auris* isolates are sensitive to echinocandins, although around 5% of isolates are reported to be resistant to this class of antifungals.(12)

Besides the high pathogenicity and the multiresistance, *C. auris* is also highly contagious. It colonizes the nose, axilla, groin and rectum, and is frequently found on inanimate surfaces and reusable equipment in healthcare facilities, which are potential sources of transmission among hospitalized patients.(13–15) It is challenging to clean contaminated surfaces and equipment, as *C. auris* can form biofilms, that are relatively insensitive to hydrogen peroxide and chlorhexidine.(16) Due to its high degree of infectivity and relative insensitivity to standard cleaning protocols, *C. auris* has caused outbreaks in various healthcare institutions, especially in intensive care settings.(13)

In 2017 whole-genome sequencing (WGS) demonstrated the presence of four different *C. auris* clades.(17) These specific geographical clades were identified as the South Asian, South American, South African, and East Asian clade. Subsequently, WGS study of *C. auris* isolates from various hospitals within the USA demonstrated the presence of all four genetically diverse clades suggesting that US patients acquired isolates and or infections from three continents.(18) This spread of *C. auris* in USA demonstrates that travel and or migration plays an important role in spreading this disease.

The identification of *C. auris* in a routine microbiology laboratory is difficult. As it is often misidentified as another *Candida* spp., like *C. haemulonii*, *C. sake*, *C. famata*, *C. lusitaniae* and *C. parapsilosis* or as *Cryptococcus laurentii* or *Rhodotorula glutinis,* the exact burden of *C. auris* outbreaks remains challenging.(19–21) A specific, rapid, accurate, reproducible and easy typing method is essential to determine the presence of a potential outbreak, however so far such method is not yet available for *C. auris*. In this study we developed a short tandem repeat (STR)-based typing for *C. auris* and used it to type a global collection of isolates.

## Results

### Selection of STR markers

Tandem repeats in the *C. auris* genome were identified using genomic information of four isolates originating from different clades. Then 23 tandem repeats of two, three, six or nine nucleotides were selected, which shared at least three repeats with an identical unit length. In order to map the 200-300 bases flanking the tandem repeats at both sides, primers were designed (**Supplementary Table S1**) and applied in PCR amplification using ten isolates from 4 known clades. PCR products were found for 22 tandem repeats and these were sequenced. One of the tandem repeats was not present in all clades, while the flanking sequence of five tandem repeats harbored deletions/insertions close to the tandem repeat, making them unsuitable for STR analysis (**Supplementary Table S1**). After excluding two repeats which showed very little variability in copy number between the isolates (**Supplementary Table S1**), a total of 14 tandem repeats remained. As this selection only included two tandem repeats with a length of six nucleotides, these were also excluded, leaving three dinucleotide, six trinucleotide and three nona-nucleotide repeats.

### *Development of* C. auris *STR typing assay and its application on a global collection of isolates*

To develop a STR typing assay for *C. auris*, primers were designed in close proximity to the tandem repeat. After testing and optimizing these primers, they were coupled to fluorescent probes. The four multiplex PCRs (M2, M3-I, M3-II and M9) were then used to genotype a *C. auris* collection of 444 isolates. All isolates were successfully typed using these multiplex panels, with the exception of isolates from Korea and Japan, which required monoplex typing of the M3-II panel. Most repeats, with the exception of the nona-nucleotide repeats, demonstrated multiple stutter peaks, due to established PCR artefacts.(22) An overview of repeat characteristics, number of alleles and discriminatory indices is shown in **Table 1**. Among 444 *C. auris* isolates, 40 different genotypes were identified containing 1 to 125 isolates (**Fig. 1A**). The genotypes clustered in 5 different groups, previously identified via WGS as the 5 different *C. auris* clades.(17, 23) These 5 clades were differentiated by at least 8 to 10 STR markers. Less variation was found within the different clades, as the maximal number of different STR markers between isolates within the South Indian clade was 7, while within the other *C. auris* clades, excluding Iran, isolates differed in maximally 3 STR markers. The total number of different alleles for the 3 markers in the M2 panel was 5, while there were respectively 6-7 and 3-4 alleles in the M3-II and M9 panel. M3-Ib and M3-Ic exhibited respectively 8 and 9 alleles, while there were 20 alleles for STR marker M3-Ia. To visualize the genotypes and the country of origin, all individual isolates are shown in a minimum spanning tree (**Fig. 1B**), which demonstrated that some genotypes were found in different countries. Remarkably, one of the South-African isolates localized in the South-American clade. Whole genome sequencing confirmed the overlap of this isolate with the South-American isolates (accession number comes here; soon available).

**Figure 1.**
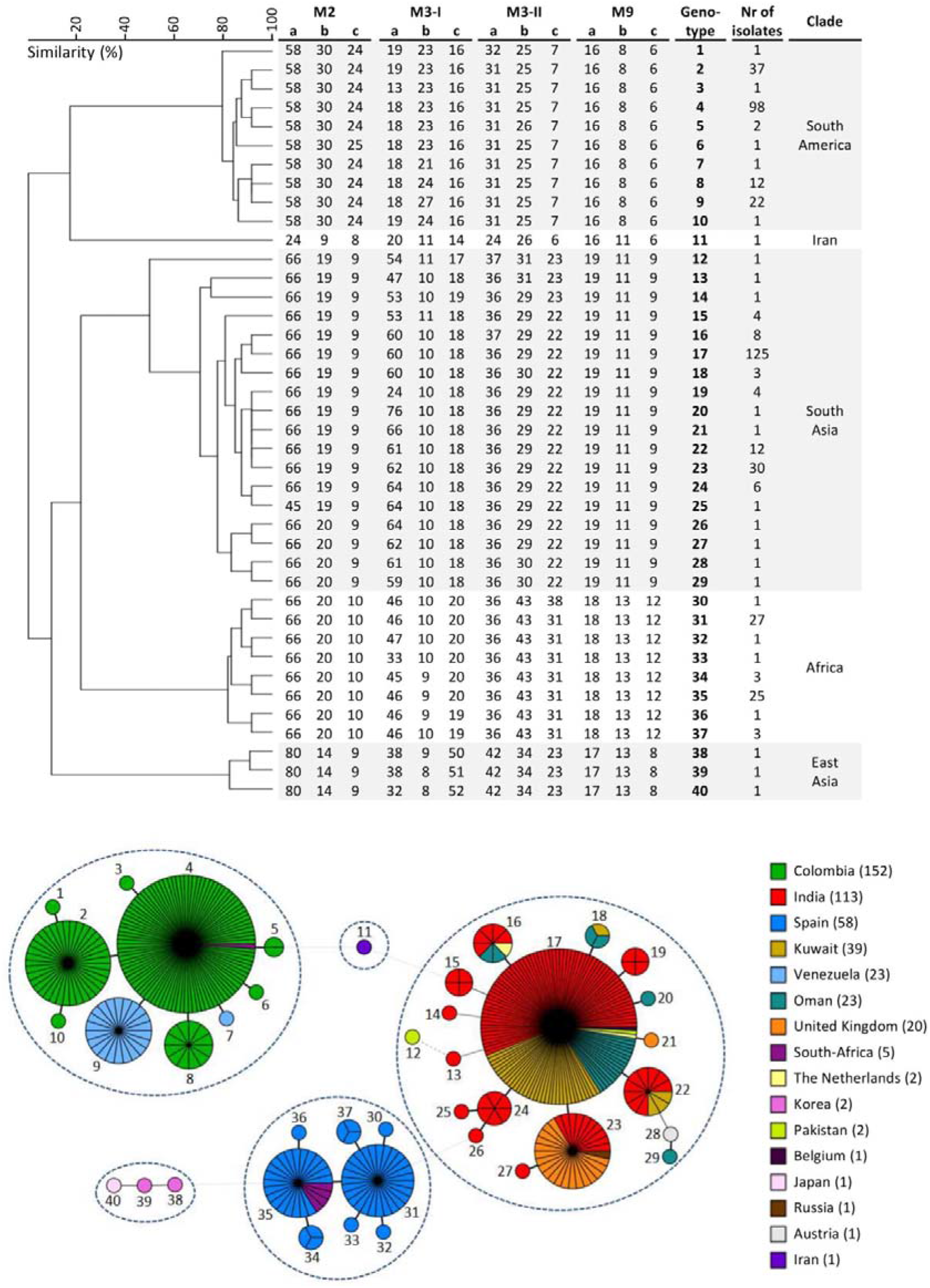
STR genotypes and minimum spanning tree of 444 *C. auris* isolates. UPGMA dendrogram of STR genotypes (upper panel) and mimimum spanning tree (lower panel) of 444 *C. auris* isolates originating from various countries. Cluster analysis showed that the different clades form distinct clusters based on the STR profiles, demarcated in the dendrogram by the grey background. The branch lengths in the minimum spanning tree (MST) indicate the similarity between isolates with thick solid lines (variation in one marker), thin solid line (variation in two markers), thin dashed lines (variation in three markers) and thin dotted lines (variation in more than 8 markers). The numbering in the MST indicates the genotype, while the number of isolates per country are described in the legend.

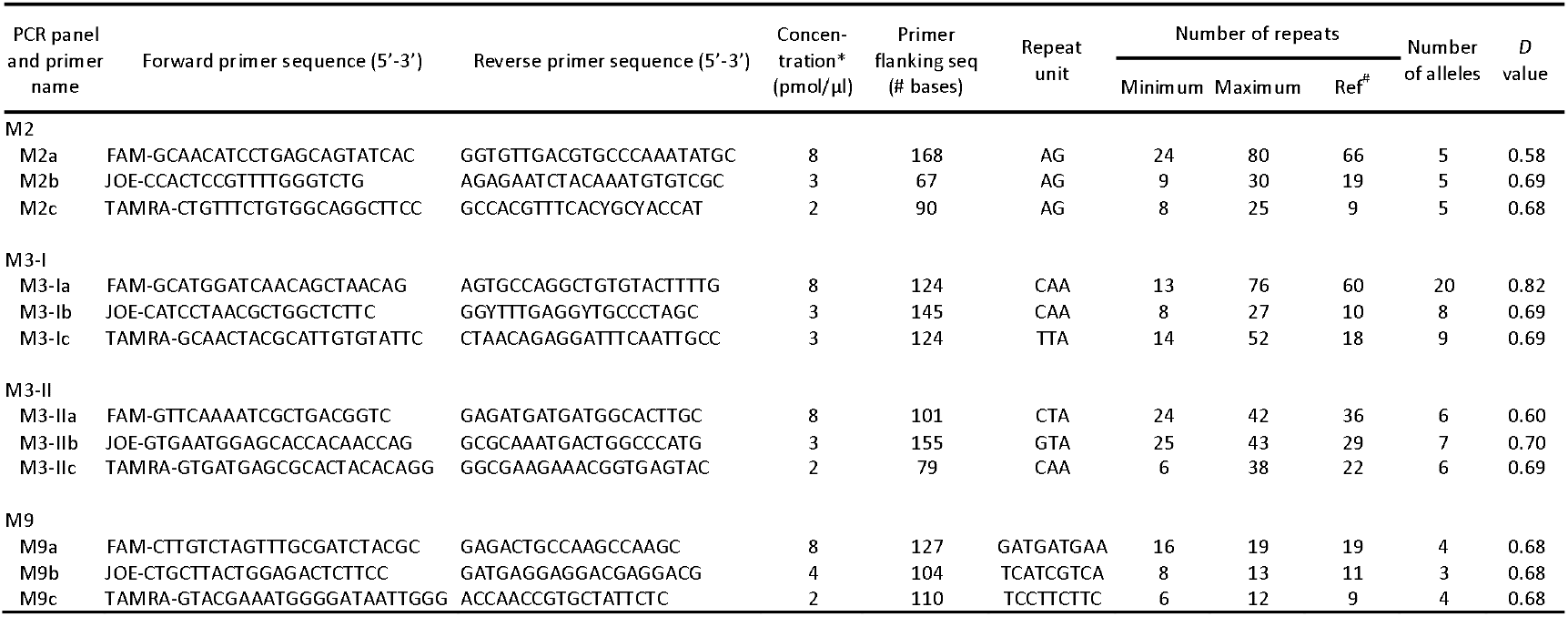

### *Reproducibility and specificity of* C. auris *STR*

To test reproducibility, isolates GMR-OM028 and VPCI 213/P/15 were independently amplified for 5 times in quadruple. STR typing demonstrated identical results for both isolates for all STR markers, demonstrating that the method is highly reproducible. In order to determine the specificity of our STR assay for *C. auris* we analyzed the following yeasts for all 12 markers: *C. haemulonii*, *C. pseudohaemulonii*, *C. duobushaemulonii*, *C. tropicalis*, *C. dubliniensis*, *C. albicans*, *C. glabrata, C. glaebosa*, *C. krusei*, *C. lusitaniae*, *C. parapsilosis*, *C. sake, Cryptococcus gattii*, *Cryptococcus albidus* and *Rhodotorula glutinis*. Products were only found with *C. duobushaemulonii* using the M3-IIb and M3-IIc marker and with *C. pseudohaemulonii* using the M2c marker, demonstrating that in general the markers are highly specific for *C. auris*.

## Discussion

The present study describes the development of a novel STR genotyping analysis for *C. auris*. This *C. auris* specific STR assay consists of 4 multiplex PCR reactions, which amplify 12 STR targets with a repeat size of 2, 3 or 9 nucleotides (panels M2, M3-I, M3-II and M9). Notably, analysis of the STR genotyping using 444 *C. auris* isolates from various geographical regions showed that this novel technique is highly reproducible and specific for *C. auris*. Furthermore, STR typing yielded highly concordant results with WGS and yielded 5 similar distinct groups, which correspond with the four well-known clades from South America, South Africa, South Asia and East Asia, and the possible new clade from Iran.^16,17,31^ The allelic variations in the 444 isolates resulted in 40 different genotypes. This relatively low number of different genotypes is partly due to the fact that >95% of our isolates originated from hospital outbreaks, leading to the inclusion of many clonal strains. Furthermore, it is known that *C. auris* only recently emerged and that there is still little variation between isolates from the same clade, as also shown by WGS.(18)

In order to get more insight in the utility of this new STR assay to type strains in an outbreak of *C. auris*, we compared its results with strains previously analyzed by WGS performed by the Centers for Disesase Control and Prevention (CDC).(17, 18) In the present study we typed 25 isolates, obtained from India, Pakistan, South-Africa and Venezuela by STR which were previously analyzed using WGS by Lockhart et al. (**Supplementary Figure S1**).(17) WGS demonstrated 4 clades, differentiated by at least tens of thousands of SNPs, while via STR analysis these 4 clades remarkably differed in at least 8 markers. Within the South-American clade, the isolates from Venezuela (n=5) were differentiated by maximal 17 SNPs, while STR profiles from these isolates were identical. A total of 4 isolates from South-Africa analyzed with both methods were differentiated by maximal 11 SNPS by WGS, and did not show any difference by STR analysis. From the South-Asian clade 14 out of 15 isolates were differentiated by ~60 SNPs by WGS, while with STR analysis these isolates demonstrated variations in marker M3-Ia and M3-Ib (genotypes 15, 17 and 22-24). Out of these 14 isolates there were 5 and 7 isolates with identical STR data (genotypes 17 and 23) with maximally 55 and 29 SNPs respectively. Interestingly, two of these isolates (B11209 with genotype 15 and B11214 with genotype 23), which originated from hospitals in Kochi in 2013 and New Delhi in 2014, India were found to be identical with WGS,(18) while STR analysis demonstrated differences in two STR markers. The fifteenth isolate, B8411, harbored ~800 SNPs as compared to the other 13 isolates,(17, 18) which corresponded with 6 different STR markers (genotype 12). Interestingly, Chow et al. demonstrated with WGS that the Japanese isolate #JCM 15448 differed in 34 and 35 SNPs compared to the two isolates KCTC17809 and KCTC17810 from South Korea (personal communication), while STR typing demonstrated differences in 2-3 markers between the Japanese and South-Korean isolates.(18) The South-Korean isolates were differentiated by 19 SNPs with WGS and one copy number in two STR markers. Altogether, isolates that differed few SNPs (<20) via WGS and are labelled as almost indistinguishable are often also not differentiated with STR analysis, while most isolates that differed in 30 or more SNPs are differentiated by STR analysis in one or more STR markers.

Implementation of a typing method in an outbreak setting requires establishing cut-off values to determine the potential relatedness of isolates. As the mutation rate between microorganisms strongly differs, such cut-off value should be determined for each microorganism separately.(24) Furthermore, methods should be standardized to determine the precise genetic variation during clonal outbreaks. This standardization is still lacking for the WGS analysis on *C. auris*. Different methods are currently used, which cannot be directly compared. This is apparent when the numbers of SNPs between identical isolates differ around 10-fold in two different labs.(18, 25) STR analysis is a standardized technique specifically developed for a single microorganism. To establish a cut-off value to determine the relatedness between isolates, we analyzed the variation between several hospital outbreaks included in this study. The London, UK 2015-2016 outbreak showed that all but one isolate exhibited a single genotype.(25) The difference of 4 copy numbers was observed in marker M3-Ia, suggesting that small variations (copy number <5) in STR marker M3-Ia may not be used to regard strains as non-related. Analysis of the outbreak in a Spanish hospital(26) showed 8 different genotypes largely originating from differences in M3-Ia marker, although also differences in three other M3 markers were found. All genotypes in Spanish outbreaks localized in the South-African clade. Due to non-availability of WGS and epidemiological information of Spanish isolates it is not possible to understand whether this population was possibly genetically heterogeneous, however the larger number of genotypes within one outbreak suggest several unique genotypes in the Spanish hospital. From the outbreaks in Colombia, we found that isolates originating from hospitals in Santa Marta, Cartagena and Medellin all exhibited genotype 4. Most isolates from the outbreaks in Popayan and Bogota exhibited a single genotype (2 and 4, respectively) although in both outbreaks there were a few single isolates with other genotypes, caused by one copy number in one marker. Finally, most isolates from Barranquilla, which originated from one hospital and were isolated between April 2015 and January 2019, clustered in two larger groups, while a few isolates exhibited a different genotype. Also, these isolates differed only in one copy number of one marker with the exception of isolate C72900 (date of isolation 28-12-2018), which exhibited 13 repeats for M3-I, while the other isolates had 18 or 19 repeats. Thus, the variation between the Colombian isolates was minimal, with exception of isolate C72900, found at the very end of the outbreak, which might be an independent introduction. Altogether, the cut-off for relatedness for the *C. auris* STR remains to be established, although small variations (copy number <5) in STR marker M3-Ia or one copy number in a M2 or other M3 STR marker should likely not be used to label strains as non-related. Our data suggests that isolates are not related with larger variation in STR markers.

The reproducibility of this STR typing assay for *C. auris* was analyzed by independent amplification of two isolates for 5 times in quadruple. As no differences were found between the typing results of WGS and STR, the novel STR assay is considered as highly reproducible. In addition to reproducibility, the specificity was also tested on different *Candida* species, and other yeasts close to *C. auris* and in the majority of targets no amplification was found. In summary we identified a STR typing assay for *C. auris* which is reliable, reproducible, specific, and cost-effective, and can be used to quickly characterize *C. auris* isolates during an outbreak.

## Materials and Methods

### Identification STR loci

Genome scaffolds of B8441-Pakistan, B11220-Japan, B11221-South Africa and B11243-Venezuela were downloaded from NCBI.(17) Scaffolds from each isolate were combined in one fasta file (http://www.bioinformatics.org/sms2/combine_fasta.html) and this fasta file was uploaded in tandem repeat finder (www.tandem.bu.edu/trf/trf.html) using the basic search option. From the resulting list of STRs, those repeats that contained insertions or deletions, exhibited <90% perfect match of the repeat sequence, did not vary in copy number between the isolates or contained repeat sequences within the potential PCR primer regions were excluded. From this list only STRs were selected from which there were at least three with an identical unit length.

### Primer design, PCR and genotyping

Primers were designed using the Tm calculator and Multiple Primer Analyzer from ThermoFisher Scientific (http://www.thermofisher.com), ordered via Eurogentec (Köln, Germany) and dissolved in stock solutions of 100 pmol/μl in 1×TE. The PCR reaction for amplification of the STR flanking regions contained 1× Fast Start Taq Polymerase buffer with MgCl_2_, 0.2 mM dNTPs, 25 pmol forward (fwd) and (rev) primer, 1 U Faststart Taq polymerase (Roche Diagnostics, Germany) and DNA and was supplemented with water to a final volume of 25 μl. A similar setup was used for the multiplex PCR reactions with 4.5 - 20 pmol fwd or rev primers. Thermocycling (Westburg, Biometra, Göttingen, Germany) was performed by using the following thermal protocol: 4 min of denaturation at 94°C, followed by 20 cycles of 15 s of denaturation at 94°C, 15 s of annealing at 66°C with a decreasing temperature of 0.5°C per cycle, and 1 min of extension at 72°C, which was then followed by 30 cycles of 15 s of denaturation at 94°C, 15 s of annealing at 56°C, and 1 min of extension at 72°C. For the multiplex PCR reactions, there was 10 min of denaturation at 95°C, followed by 30 cycles of 30 s of denaturation at 95°C, 30 s of annealing at 60°C, and 1 min of extension at 72°C. Before the reaction mixtures were cooled to room temperature, an additional incubation for 10 min at 72°C was performed. For DNA sequencing, the product was purified according to the Ampliclean method (NimaGen, Nijmegen, Netherlands), and the sequencing PCR was performed using 0.5 μl BrilliantDye premix, 1.75 μl BrilliantDye 5× sequencing buffer (NimaGen), 5 pmol fwd or rev primer, 5.75 μl water and 1 μl DNA with the following thermal protocol: 45 s of denaturation at 96°C, followed by 35 cycles of 10 s of denaturation at 96°C, 5 s of annealing at 50°C, and 2 min of extension at 60°C. After D-Pure purification (Nimagen, Nijmegen, Netherlands) sequencing was performed using the 3500XL genetic analyzer (Applied Biosystems, Foster City, CA, USA) and analyzed in Bionumerics 7.6.1 (Applied Maths, Kortrijk, Belgium). For the STR analysis, samples were diluted 1:1000 and 10 μl of the diluent, together with 0.12 μl Orange 600 DNA Size Standard (Nimagen), was boiled for 1 min at 95°C, and analyzed according to the manufacturer’s recommendations on an automatic sequencer ABI 3500XL Genetic Analyzer (Applied Biosystems).

### Whole genome sequencing

Genomic libraries were prepared and sequenced with Illumina technology (Illumina, San Diego, USA) with 2 × 150 bp paired-end read mode at Eurofins Genomics (Ebersberg, Germany). Seventy-four *C. auris* WGS sequences from NCBI were added to the analysis. FastQC and PRINSEQ was used to assess quality of read data and perform read filtering. Read data were aligned against a publically available genome sequenced on PacBio RS II using BWA. SNP variants were identified using SAMtools and filtered using the publically available SNP analysis pipeline NASP to remove positions that had less than 10x coverage, less than 90% variant allele calls, or that were identified by Nucmer as being within duplicated regions in the reference. Phylogenetic analysis and bootstrapping with 1000 iterations were performed on SNP matrices using RAxML.

### Isolates

For the STR analysis 444 *C. auris* isolates from Austria(27), Belgium(28), Colombia (hospitals in Barranquilla, Bogota, Cartagena, Medellin, Popayan and Santa Maria), India(29), Iran(30), Japan(29), Korea(29), Kuwait(31), Oman(32), Pakistan(17), Russia(33), South-Africa(29), Spain(26), The Netherlands(34), United Kingdom(35) and Venezuela(6) were used. For a complete overview of isolates see **Supplementary Table S2**. Isolates were stored at −80°C, according to standard procedures. Species identification via sequencing and/or MALDI-TOF MS was carried out as described previously.(29)

The specificity of the STR was tested on the spectra of yeast isolates including those that are previously known to yield misidentification of *C. auris* by commercial biochemical methods. The following isolates *C. haemulonii* (CBS5149T, CBS7802, CBS7801 and CBS5150), *C. pseudohaemulonii (JCM12453T*, KCTC1787, CBS10004T), *C. duobushaemulonii* (CBS7796, CBS7800, CBS7799, CBS9754), *C. tropicalis* (ATCC750), *C. dubliniensis* (CBS8500), *C. albicans* (ATCC10231), *C. glaebosa* (clinical isolate), *C. krusei* (ATCC6258), *C. glabrata* (ATCC2175), *C. lusitaniae* (CWZ ID 10-11-02-05), *C*. *parapsilosis* (CWZ ID 10-06-05-86), *C. sake* (clinical isolate), *Cryptococcus gattii* (CBS12652), *Cryptococcus albidus* (clinical isolate) and *Rhodotorula glutinis* (CWZ ID 10-06-05-66) were tested.

### Culture and DNA isolation

Isolates were grown on Sabouraud agar plates at 35°C. For DNA sequencing, strains were resuspended in a vial with 400 μl MagNA Pure Bacteria Lysis Buffer and MagNA Lyser Green Beads beads and mechanically lysed for 30 sec at 6,500 rpm using the MagNA Lyser (all Roche Diagnostics GmbH, Mannheim, Germany). Subsequently, DNA was extracted and purified with the MagNA Pure LC instrument and the MagNA Pure DNA isolation kit III (Roche Diagnostics), according to the recommendations of the manufacturer. For STR analysis, strains were resuspended in 50 μl physiological salt and after addition of 200 U of lyticase (Sigma-Aldrich, St. Louis, MO, USA) and incubation for 5 min at 37 °C, 450 μl physiological salt was added. The sample was then incubated for 15 min at 100 °C, and cooled down to room temperature.

### Data analysis and discriminatory power

Copy number of the twelve markers of all isolates were determined using GeneMapper Software 5 (Applied Biosystems). Relatedness between isolates was analyzed using BioNumerics, version 7.6.1, software (Applied Maths) via the unweighted pair group method with arithmetic averages using the multistate categorical similarity coefficient. All markers were given an equal weight. The discriminatory power of the STR assay was determined using the Simpson index of diversity (D) as described previously.(22) A ‘D’ value of 1.0 indicates that according to the used typing method all isolates have a different genotype, while a ‘D’ value of 0 indicates all isolates to be identical.

## Acknowledgements

This research was financially supported by the Canisius Wilhelmina Hospital. The authors thank Dr. Nancy Chow for help with WGS analysis.

---

**Supplementary Figure S1** Comparison genotypic difference of *C. auris* isolates via STR and WGS analysis. Lockhart et al provided detailed information regarding the precise number of SNPs differentiating 47 *C. auris* isolates via WGS (left panel), which was compared with the STR genotypes of isolates that were included in the present study (right panel). The maximal differences in SNPs between isolates with genotype 23, 17, 35 and 9 was respectively 29, 55, 11 and 17. Gt, genotype.

