## Supplementary Figure S1 for "Development of *Candida auris* microsatellite typing and its application on a global collection of isolates"

**Supplementary Figures -** Supplementary Figure S1

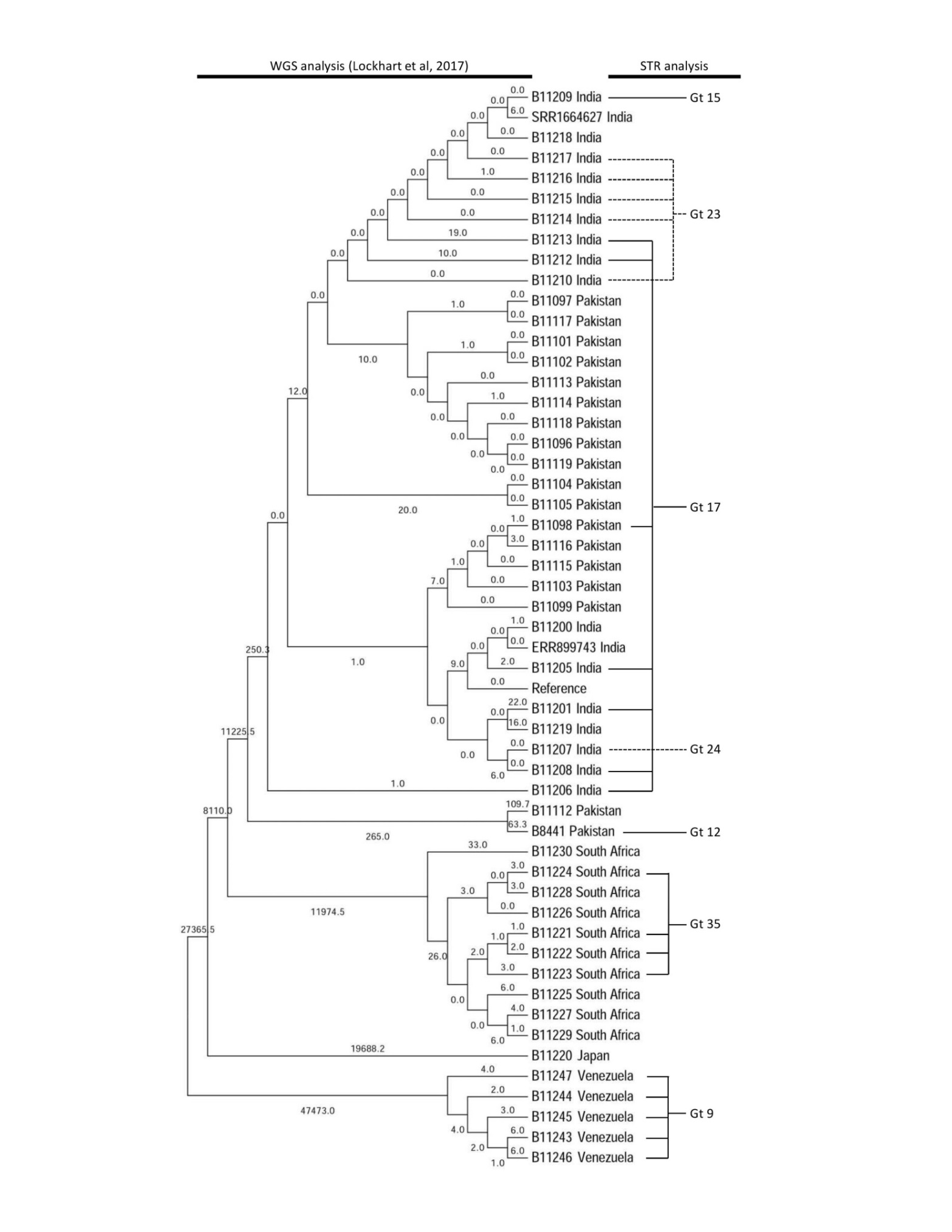

**Supplementary Figure S1** Comparison genotypic difference of *C. auris* isolates via STR and WGS analysis. Lockhart et al provided detailed information regarding the precise number of SNPs differentiating 47 *C. auris* isolates via WGS (left panel), which was compared with the STR genotypes of isolates that were included in the present study (right panel). The maximal differences in SNPs between isolates with genotype 23, 17, 35 and 9 was respectively 29, 55, 11 and 17. Gt, genotype.
