## Supplementary Table S1 for "Development of *Candida auris* microsatellite typing and its application on a global collection of isolates"

| STR code | Repeat | Primer | Sequence (5’-3’) | STR present in all isolates tested | Absence of deletions or insertions close to STR | Variability in copy number | Locus sequence |
| --- | --- | --- | --- | --- | --- | --- | --- |
| 2-1 | AG | 2-1Fwd  2-1Rev | CAACACTGCATGGACCAAAGCG  GACTCATTCGCACTTGCCTCG | Yes | Yes | Yes |  |
| 2-2 | GA | 2-2Fwd  2-2Rev | GAGGTATGTTCTTGCATCACCAC  GAGCTATCTCGTGCTTCTTACC | No | - | - | - |
| 2-3 | AG | 2-3Fwd  2-3Rev | CTTGAGATCCAGTGACAATCT  CACCACCATGTAACTAGCG | Yes | Yes | Yes |  |
| 2-4 | CT | 2-4Fwd*  2-4Fwd2*  2-4Rev*  2-4Rev2*  2-4Rev3*  2-4Rev4*  2-4Rev5* | GGTTGCATGTGGACCAACTTC  GCTTCTCCTTATAGGCCGCTTAC  CCATTCTTTACTGTTCCGAATCG  GAGCCAATAACTTTACCATTC  CTTTACCATTCTTTACTGTTCCG  CCTTCAATGATCTAATGACGGC  GCAACAACGTAAGCGGACTTCG | - | - | - | - |
| 2-5 | AG | 2-5Fwd  2-5Rev | CCACACTTCTGCAGGAAACGTGG  GAGCTGTGAGGCCAAGATCCCTC | Yes | Yes | Yes |  |
| 3-1 | CAA | 3-1Fwd  3-1Rev | GATCTCGACTTCGACCTGCTCTTG  CTGAGGGTTCGAGGAGAATTGCTG | Yes | Yes | Yes |  |
| 3-2 | CAA | 3-2Fwd  3-2Rev | CTACCGAACAATCCACTGCCTTC  CTTGCTCTCCAGAACTTGCTGG | Yes | Yes | Yes |  |
| 3-3 | TTA | 3-3Fwd  3-3Rev | GGATCTCCACTTGGTAGTCGTCG  GCGACAAGTTCAAGCGTGAGTAG | Yes | Yes | Yes |  |
| 3-4 | CTA | 3-4Fwd*  3-4Rev*  3-4Fwd2  3-4Rev2 | GTCGTGCCACACTGCTTCTTC  CGTCATTCGCCAAGAACCTAG  CGAACTGTTCTTCCTCACTACC  GCACCTATCGAAGGTATTGACG | Yes | Yes | Yes |  |
| 3-5 | GTA | 3-5Fwd  3-5Rev | CGTACAGCAGCCTTTAGCATCC  GCGTGCTTGGTGCATCCATTG | Yes | Yes | Yes |  |
| 3-6 | CAA | 3-6Fwd  3-6Rev | GACAAACCAGGCGTATCGC  CAGCTGGCCTGATAAGAGTGGC | Yes | No | - | - |
| 3-7 | TGT | 3-7Fwd  3-7Rev | GCTTCTTCTGGCTCTGGCTCTG  GGAGAACTTACCCACGATCCCAG | Yes | No | - | - |
| 3-8 | CAG | 3-8Fwd  3-8Rev | CTACCTCAGCAGGAGAGTATCC  CAGAAGCTGCCTGCTCGTTG | Yes | Yes | No | - |
| 3-9 | TGT | 3-9Fwd  3-9Rev | GTACAGCTGTCGCTGATTC  CCGTGAAGATACCTGATACCTCG | Yes | Yes | No | - |
| 3-10 | CAA | 3-10Fwd  3-10Rev | CGACGTCTTGACCAAGGGTC  CACTATGCAACTGCGACAGC | Yes | Yes | Yes |  |
| 6-1 | ATACAT | 6-1Fwd  6-1Rev | CCGATGCTTTGTAGTGATGCCAT  GAGGCATGTTCTCTATGCCACAG | Yes | No | - | - |
| 6-2 | CAGAAC | 6-2Fwd  6-2Rev | CTAGTAGCAAGCCCACCAAGGTC  CTTCTCGACGGTGATACGCTTC | Yes | Yes | Yes | - |
| 6-3 | AGCTCT | 6-3Fwd  6-3Rev | CCACTCTGTATGGTCCAAGGC  CTCACCAGTAGCCTCTGGAGTTG | Yes | Yes | Yes | - |
| 9-1 | GATGATGAA | 9-1Fwd*  9-1Rev*  9-1Fwd2  9-1Rev2 | GTAGCGTGCTTAGAGCTTCG  CAAGCGAACTCAACTCTATGCC  CATACTGCTTATTGGGCCAGAG  GTTGTGATTCCTCGATAGCAG | Yes | Yes | Yes |  |
| 9-2 | GGTCCAAGC | 9-2Fwd  9-2Rev | GACTCTCACACTGGTCCATCC  GGTATCAACGTACATTAGGCG | Yes | No | - | - |
| 9-3 | TCATCGTCA | 9-3Fwd*  9-3Rev*  9-3Fwd2  9-3Rev2 | GGACGTGTTACCTCTTCTTGAACTG  CCTGCAATAGCTCCTGTGGGAC  CTACATCACTTCCAACTACAGC  GTACACACATCCCATGACAGC | Yes | Yes | Yes |  |
| 9-4 | GGACCTAGT | 9-4Fwd  9-4Rev | CAACTGAGTCAACCACCTCAGGG  GTTGCAAGTCTCACAGCTCC | Yes | No | - | - |
| 9-5 | TCCTTCTTC | 9-5Fwd  9-5Rev | GCTCTTCTTCGTACCTCTAAGC  CATCATTGGTCGGTCAGGCTAC | Yes | Yes | Yes |  |

* PCR was not successful for one or more isolates.
