## Supplementary Table S2 for "Development of *Candida auris* microsatellite typing and its application on a global collection of isolates"

| **Geno-type** | **Isolate ID** | **Isolate ID-Alt.1** | **Isolate ID-Alt.2** | **Country** | **City** |
| --- | --- | --- | --- | --- | --- |
| 1 | GMR-OM473 |  |  | Colombia | Popayan |
| 2 | C52710-20 |  |  | Colombia | Barranquilla |
| 2 | COL 001 |  |  | Colombia | Bogota |
| 2 | COL 446 |  |  | Colombia | Bogota |
| 2 | GMR-OM043 |  |  | Colombia | Popayan |
| 2 | GMR-OM045 |  |  | Colombia | Popayan |
| 2 | GMR-OM047 |  |  | Colombia | Popayan |
| 2 | GMR-OM048 |  |  | Colombia | Popayan |
| 2 | GMR-OM050 |  |  | Colombia | Popayan |
| 2 | GMR-OM070 |  |  | Colombia | Popayan |
| 2 | GMR-OM072 |  |  | Colombia | Popayan |
| 2 | GMR-OM073 |  |  | Colombia | Popayan |
| 2 | GMR-OM077 |  |  | Colombia | Popayan |
| 2 | GMR-OM078 |  |  | Colombia | Popayan |
| 2 | GMR-OM081 |  |  | Colombia | Popayan |
| 2 | GMR-OM082 |  |  | Colombia | Popayan |
| 2 | GMR-OM085 |  |  | Colombia | Popayan |
| 2 | GMR-OM113 |  |  | Colombia | Popayan |
| 2 | GMR-OM115 |  |  | Colombia | Popayan |
| 2 | GMR-OM118 |  |  | Colombia | Popayan |
| 2 | GMR-OM193 |  |  | Colombia | Popayan |
| 2 | GMR-OM295 |  |  | Colombia | Popayan |
| 2 | GMR-OM452 |  |  | Colombia | Popayan |
| 2 | GMR-OM454 |  |  | Colombia | Popayan |
| 2 | GMR-OM458 |  |  | Colombia | Popayan |
| 2 | GMR-OM480 |  |  | Colombia | Popayan |
| 2 | GMR-OM516 |  |  | Colombia | Popayan |
| 2 | GMR-OM519 |  |  | Colombia | Popayan |
| 2 | GMR-OM524 |  |  | Colombia | Popayan |
| 2 | GMR-OM525 |  |  | Colombia | Popayan |
| 2 | GMR-OM527 |  |  | Colombia | Popayan |
| 2 | GMR-OM532 |  |  | Colombia | Popayan |
| 2 | GMR-OM743 |  |  | Colombia | Popayan |
| 2 | GMR-OM748 |  |  | Colombia | Popayan |
| 2 | GMR-OM753 |  |  | Colombia | Popayan |
| 2 | GMR-OM775 |  |  | Colombia | Popayan |
| 2 | GMR-OM781 |  |  | Colombia | Popayan |
| 2 | GMR-OM828 |  |  | Colombia | Popayan |
| 3 | C72900 |  |  | Colombia | Barranquilla |
| 4 | 1976339 |  |  | Colombia | Barranquilla |
| 4 | A106871-47 |  |  | Colombia | Barranquilla |
| 4 | A110602-27 |  |  | Colombia | Barranquilla |
| 4 | A145737 |  |  | Colombia | Barranquilla |
| 4 | C45329-14 |  |  | Colombia | Barranquilla |
| 4 | C46052-13 |  |  | Colombia | Barranquilla |
| 4 | C46305-04 |  |  | Colombia | Barranquilla |
| 4 | C47103-43 |  |  | Colombia | Barranquilla |
| 4 | C47129-34 |  |  | Colombia | Barranquilla |
| 4 | C47453-32 |  |  | Colombia | Barranquilla |
| 4 | C47753 |  |  | Colombia | Barranquilla |
| 4 | C48117-22 |  |  | Colombia | Barranquilla |
| 4 | C48321-02 |  |  | Colombia | Barranquilla |
| 4 | C48374-12 |  |  | Colombia | Barranquilla |
| 4 | C48406-05 |  |  | Colombia | Barranquilla |
| 4 | C48477 |  |  | Colombia | Barranquilla |
| 4 | C48522-11 |  |  | Colombia | Barranquilla |
| 4 | C48621-38 |  |  | Colombia | Barranquilla |
| 4 | C49302-40 |  |  | Colombia | Barranquilla |
| 4 | C52043-49 |  |  | Colombia | Barranquilla |
| 4 | C52710-21 |  |  | Colombia | Barranquilla |
| 4 | C55038-53 |  |  | Colombia | Barranquilla |
| 4 | C56029-45 |  |  | Colombia | Barranquilla |
| 4 | C56237-48 |  |  | Colombia | Barranquilla |
| 4 | C56305-55 |  |  | Colombia | Barranquilla |
| 4 | C57942-25 |  |  | Colombia | Barranquilla |
| 4 | C59378-23 |  |  | Colombia | Barranquilla |
| 4 | C59545-41 |  |  | Colombia | Barranquilla |
| 4 | C59629-54 |  |  | Colombia | Barranquilla |
| 4 | C60318-26 |  |  | Colombia | Barranquilla |
| 4 | C60470-17 |  |  | Colombia | Barranquilla |
| 4 | C60912-35 |  |  | Colombia | Barranquilla |
| 4 | C60920-36 |  |  | Colombia | Barranquilla |
| 4 | C61398-07 |  |  | Colombia | Barranquilla |
| 4 | C61405-01 |  |  | Colombia | Barranquilla |
| 4 | C61697-08 |  |  | Colombia | Barranquilla |
| 4 | C62320-03 |  |  | Colombia | Barranquilla |
| 4 | C62530 |  |  | Colombia | Barranquilla |
| 4 | C62651 |  |  | Colombia | Barranquilla |
| 4 | C62959 |  |  | Colombia | Barranquilla |
| 4 | C63037 |  |  | Colombia | Barranquilla |
| 4 | C63261 |  |  | Colombia | Barranquilla |
| 4 | C63476 |  |  | Colombia | Barranquilla |
| 4 | C71792 |  |  | Colombia | Barranquilla |
| 4 | C72538 |  |  | Colombia | Barranquilla |
| 4 | C72764 |  |  | Colombia | Barranquilla |
| 4 | D26413-28 |  |  | Colombia | Barranquilla |
| 4 | D29575-44 |  |  | Colombia | Barranquilla |
| 4 | D32219 |  |  | Colombia | Barranquilla |
| 4 | G8201 |  |  | Colombia | Barranquilla |
| 4 | 386 |  |  | Colombia | Bogota |
| 4 | 445 |  |  | Colombia | Bogota |
| 4 | COL 1076 |  |  | Colombia | Bogota |
| 4 | COL 1080 |  |  | Colombia | Bogota |
| 4 | COL 885 |  |  | Colombia | Bogota |
| 4 | COL 941 |  |  | Colombia | Bogota |
| 4 | COL 990 |  |  | Colombia | Bogota |
| 4 | 12272906 |  |  | Colombia | Cartagena |
| 4 | 11106T |  |  | Colombia | Cartagena |
| 4 | JLIB |  |  | Colombia | Cartagena |
| 4 | 2001051 |  |  | Colombia | Medellin |
| 4 | 3597150 |  |  | Colombia | Medellin |
| 4 | 3597150 |  |  | Colombia | Medellin |
| 4 | 3620098 |  |  | Colombia | Medellin |
| 4 | 3636615 |  |  | Colombia | Medellin |
| 4 | 10402302 |  |  | Colombia | Medellin |
| 4 | 10500471 |  |  | Colombia | Medellin |
| 4 | 11902111 |  |  | Colombia | Medellin |
| 4 | 12902191 |  |  | Colombia | Medellin |
| 4 | 20201961 |  |  | Colombia | Medellin |
| 4 | 21000622 |  |  | Colombia | Medellin |
| 4 | 21401141 |  |  | Colombia | Medellin |
| 4 | 121000761 |  |  | Colombia | Medellin |
| 4 | 121001051 |  |  | Colombia | Medellin |
| 4 | 121001401 |  |  | Colombia | Medellin |
| 4 | 122101981 |  |  | Colombia | Medellin |
| 4 | 122801101 |  |  | Colombia | Medellin |
| 4 | 122801141 |  |  | Colombia | Medellin |
| 4 | 122900941 |  |  | Colombia | Medellin |
| 4 | 03388273-41 |  |  | Colombia | Medellin |
| 4 | 03531221-28 |  |  | Colombia | Medellin |
| 4 | 03531221-29 |  |  | Colombia | Medellin |
| 4 | 03531221-39 |  |  | Colombia | Medellin |
| 4 | 03588820 |  |  | Colombia | Medellin |
| 4 | 121002511 (90) |  |  | Colombia | Medellin |
| 4 | 121002511 (93) |  |  | Colombia | Medellin |
| 4 | 122801121 2x |  |  | Colombia | Medellin |
| 4 | 21813229-012201971 |  |  | Colombia | Medellin |
| 4 | L12097-012816941 |  |  | Colombia | Medellin |
| 4 | GMR-OM028 |  |  | Colombia | Popayan |
| 4 | GMR-OM538 |  |  | Colombia | Popayan |
| 4 | 2274737-42 |  |  | Colombia | Santa Marta |
| 4 | 2276903-51 |  |  | Colombia | Santa Marta |
| 4 | 2321573-37 |  |  | Colombia | Santa Marta |
| 4 | 2323170-18 |  |  | Colombia | Santa Marta |
| 4 | 2324866-46 |  |  | Colombia | Santa Marta |
| 4 | 2326512-39 |  |  | Colombia | Santa Marta |
| 4 | MOL353 |  |  | SA |  |
| 5 | C47129-31 |  |  | Colombia | Barranquilla |
| 5 | F23720-29 |  |  | Colombia | Barranquilla |
| 6 | COL 880 |  |  | Colombia | Bogota |
| 7 | 22 |  | 10-08-01-15 | Venezuela |  |
| 8 | C45563-52 |  |  | Colombia | Barranquilla |
| 8 | C46772-19 |  |  | Colombia | Barranquilla |
| 8 | C48602-33 |  |  | Colombia | Barranquilla |
| 8 | C54019-50 |  |  | Colombia | Barranquilla |
| 8 | C57272-10 |  |  | Colombia | Barranquilla |
| 8 | C57506-24 |  |  | Colombia | Barranquilla |
| 8 | C58960-15 |  |  | Colombia | Barranquilla |
| 8 | C59378-30 |  |  | Colombia | Barranquilla |
| 8 | C61797-06 |  |  | Colombia | Barranquilla |
| 8 | C62035-09 |  |  | Colombia | Barranquilla |
| 8 | C71269 |  |  | Colombia | Barranquilla |
| 8 | C72508 |  |  | Colombia | Barranquilla |
| 9 | 1 |  | 10-08-01-01 | Venezuela |  |
| 9 | 2 |  | 10-08-01-02 | Venezuela |  |
| 9 | 4 |  | 10-08-01-03 | Venezuela |  |
| 9 | 5 |  | 10-08-01-04 | Venezuela |  |
| 9 | 6 |  | 10-08-01-05 | Venezuela |  |
| 9 | 7 |  | 10-08-01-06 | Venezuela |  |
| 9 | 13 |  | 10-08-01-07 | Venezuela |  |
| 9 | 15 |  | 10-08-01-08 | Venezuela |  |
| 9 | 16 |  | 10-08-01-09 | Venezuela |  |
| 9 | 17 |  | 10-08-01-10 | Venezuela |  |
| 9 | 18 |  | 10-08-01-11 | Venezuela |  |
| 9 | 19 |  | 10-08-01-12 | Venezuela |  |
| 9 | 20 |  | 10-08-01-13 | Venezuela |  |
| 9 | 21 |  | 10-08-01-14 | Venezuela |  |
| 9 | 23 |  | 10-08-01-16 | Venezuela |  |
| 9 | 24 |  | 10-08-01-17 | Venezuela |  |
| 9 | 25 |  | 10-08-01-18 | Venezuela |  |
| 9 | 129018 | B11245 | 10-05-15-22 | Venezuela |  |
| 9 | 129495 | B11247 | 10-05-15-19 | Venezuela |  |
| 9 | 129628 | B11243 | 10-05-15-20 | Venezuela |  |
| 9 | 201621 | B11244 | 10-05-15-21 | Venezuela |  |
| 9 | 1216950 | B11246 | 10-05-15-23 | Venezuela |  |
| 10 | GMR-OM830 |  |  | Colombia | Popayan |
| 11 | IFRC 2087 |  |  | Iran |  |
| 12 | CDC 387 | B8441 |  | Pakistan |  |
| 13 | CDC 382 |  |  | India |  |
| 14 | VPCI 960/P/17(1) |  |  | India |  |
| 15 | VPCI 1130/P/13 | B11209 |  | India |  |
| 15 | VPCI 1131/P/13 |  |  | India |  |
| 15 | VPCI 249/P/14 |  |  | India |  |
| 15 | VPCI 260/P/14 |  |  | India |  |
| 16 | VPCI 1560/P/17 |  |  | India |  |
| 16 | VPCI 1757/P/18 |  |  | India |  |
| 16 | VPCI 211/P/15 |  |  | India |  |
| 16 | VPCI 431/P/15 |  |  | India |  |
| 16 | VPCI 641/P/15 |  |  | India |  |
| 16 | 14 |  |  | Oman |  |
| 16 | 316015109 |  |  | Oman |  |
| 16 | Amsterdam |  |  | NL |  |
| 17 | 05-299 |  |  | Belgium |  |
| 17 | CDC 390 |  |  | India |  |
| 17 | VPCI 1030/P/17 |  |  | India |  |
| 17 | VPCI 105/P/14 | B11212 |  | India |  |
| 17 | VPCI 106/P/14 |  |  | India |  |
| 17 | VPCI 107/P/14 | B11213 |  | India |  |
| 17 | VPCI 1182/P/15 |  |  | India |  |
| 17 | VPCI 1245/P/15 |  |  | India |  |
| 17 | VPCI 1280/P/16 |  |  | India |  |
| 17 | VPCI 1306/P/16 |  |  | India |  |
| 17 | VPCI 1556/P/17 |  |  | India |  |
| 17 | VPCI 1576/P/17 |  |  | India |  |
| 17 | VPCI 1580/P/17 |  |  | India |  |
| 17 | VPCI 1593/P/17 |  |  | India |  |
| 17 | VPCI 1771/P/18 |  |  | India |  |
| 17 | VPCI 1875/P/18 |  |  | India |  |
| 17 | VPCI 1882/P/18 |  |  | India |  |
| 17 | VPCI 2439/P/16 |  |  | India |  |
| 17 | VPCI 2442/P/16 |  |  | India |  |
| 17 | VPCI 245/P/14 |  |  | India |  |
| 17 | VPCI 2451/P/16 |  |  | India |  |
| 17 | VPCI 247/P/14 |  |  | India |  |
| 17 | VPCI 261/P/14 |  |  | India |  |
| 17 | VPCI 263/P/14 |  |  | India |  |
| 17 | VPCI 266/P/14 |  |  | India |  |
| 17 | VPCI 271/P/14 |  |  | India |  |
| 17 | VPCI 458/P/17 |  |  | India |  |
| 17 | VPCI 463/P/14 |  |  | India |  |
| 17 | VPCI 464/P/14 |  |  | India |  |
| 17 | VPCI 468/P/14 |  |  | India |  |
| 17 | VPCI 469/P/14 |  |  | India |  |
| 17 | VPCI 470/P/14 |  |  | India |  |
| 17 | VPCI 471A/P/14 |  |  | India |  |
| 17 | VPCI 473/P/14 |  |  | India |  |
| 17 | VPCI 474/P/14 |  |  | India |  |
| 17 | VPCI 475/P/13 |  |  | India |  |
| 17 | VPCI 476/P/13 |  |  | India |  |
| 17 | VPCI 477/P/14 |  |  | India |  |
| 17 | VPCI 478/P/13 | B11205 |  | India |  |
| 17 | VPCI 479/P/13 |  |  | India |  |
| 17 | VPCI 480/P/13 | B11206 |  | India |  |
| 17 | VPCI 483/P/13 | B11208 |  | India |  |
| 17 | VPCI 484/P/13 |  |  | India |  |
| 17 | VPCI 484/P/13 |  |  | India |  |
| 17 | VPCI 507/P/14 |  |  | India |  |
| 17 | VPCI 507/P/14 |  |  | India |  |
| 17 | VPCI 508/P/14 |  |  | India |  |
| 17 | VPCI 509/P/15 |  |  | India |  |
| 17 | VPCI 524/P/15 |  |  | India |  |
| 17 | VPCI 539/P/15 |  |  | India |  |
| 17 | VPCI 588/P/16 |  |  | India |  |
| 17 | VPCI 603/P/16 |  |  | India |  |
| 17 | VPCI 650/P/16 |  |  | India |  |
| 17 | VPCI 652/P/16 |  |  | India |  |
| 17 | VPCI 659/P/16 |  |  | India |  |
| 17 | VPCI 669/P/12 |  |  | India |  |
| 17 | VPCI 670/P/12 |  |  | India |  |
| 17 | VPCI 671/P/12 |  |  | India |  |
| 17 | VPCI 672/P/12 |  |  | India |  |
| 17 | VPCI 673/P/12 |  |  | India |  |
| 17 | VPCI 683/P/12 | B11201 |  | India |  |
| 17 | VPCI 692/P/12 |  |  | India |  |
| 17 | VPCI 708/P/12 |  |  | India |  |
| 17 | VPCI 709/P/12 |  |  | India |  |
| 17 | VPCI 711/P/12 |  |  | India |  |
| 17 | VPCI 712/P/12 |  |  | India |  |
| 17 | VPCI 743/P/16 |  |  | India |  |
| 17 | VPCI 758/P/15 |  |  | India |  |
| 17 | VPCI 783/P/17 |  |  | India |  |
| 17 | VPCI 791/P/17 |  |  | India |  |
| 17 | VPCI 985/P/16 |  |  | India |  |
| 17 | 1079/16 |  |  | Kuwait |  |
| 17 | 1133/16 |  |  | Kuwait |  |
| 17 | 1501/16 |  |  | Kuwait |  |
| 17 | 1548/15 |  |  | Kuwait |  |
| 17 | 1770/16 |  |  | Kuwait |  |
| 17 | 1856/17 |  |  | Kuwait |  |
| 17 | 1994/17 |  |  | Kuwait |  |
| 17 | 2040/17 |  |  | Kuwait |  |
| 17 | 2074/17 |  |  | Kuwait |  |
| 17 | 2112/16 |  |  | Kuwait |  |
| 17 | 2140/16 |  |  | Kuwait |  |
| 17 | 2233/16 |  |  | Kuwait |  |
| 17 | 2260/17 |  |  | Kuwait |  |
| 17 | 2316/16 |  |  | Kuwait |  |
| 17 | 2501/17 |  |  | Kuwait |  |
| 17 | 2611/17 |  |  | Kuwait |  |
| 17 | 2690/15 |  |  | Kuwait |  |
| 17 | 2762/15 |  |  | Kuwait |  |
| 17 | 2857/14 |  |  | Kuwait |  |
| 17 | 2864/15 |  |  | Kuwait |  |
| 17 | 2946/15 |  |  | Kuwait |  |
| 17 | 348/15 |  |  | Kuwait |  |
| 17 | 3626/15 |  |  | Kuwait |  |
| 17 | 3716/14 |  |  | Kuwait |  |
| 17 | 3758/15 |  |  | Kuwait |  |
| 17 | 3824/15 |  |  | Kuwait |  |
| 17 | 3833/15 |  |  | Kuwait |  |
| 17 | 3849/15 |  |  | Kuwait |  |
| 17 | 3910/15 |  |  | Kuwait |  |
| 17 | 39825/15 |  |  | Kuwait |  |
| 17 | 541/16 |  |  | Kuwait |  |
| 17 | 61/16 |  |  | Kuwait |  |
| 17 | 636/15 |  |  | Kuwait |  |
| 17 | 809/16 |  |  | Kuwait |  |
| 17 | 850/16 |  |  | Kuwait |  |
| 17 | 12 |  |  | Oman |  |
| 17 | 317016400 |  |  | Oman |  |
| 17 | 317052804 |  |  | Oman |  |
| 17 | 318005583 |  |  | Oman |  |
| 17 | 318014160 |  |  | Oman |  |
| 17 | 318014517 |  |  | Oman |  |
| 17 | CHL 1692 |  |  | Oman |  |
| 17 | CHL 1874 |  |  | Oman |  |
| 17 | CHL 1906 |  |  | Oman |  |
| 17 | CHL 1940 |  |  | Oman |  |
| 17 | CHL 1941 |  |  | Oman |  |
| 17 | CHL 2170 |  |  | Oman |  |
| 17 | CHL 2830 |  |  | Oman |  |
| 17 | CHL 2982 |  |  | Oman |  |
| 17 | CHL 3182 |  |  | Oman |  |
| 17 | CHL 3234 |  |  | Oman |  |
| 17 | CHL 3236 |  |  | Oman |  |
| 17 | CDC 388 | B11098 |  | Pakistan |  |
| 17 | 8032139565 |  |  | NL |  |
| 18 | 291/15 |  |  | Kuwait |  |
| 18 | 318015306 |  |  | Oman |  |
| 18 | CHL 2795 |  |  | Oman |  |
| 19 | CDC 389 |  |  | India |  |
| 19 | VPCI 471/P/13 |  |  | India |  |
| 19 | VPCI 472/P/13 |  |  | India |  |
| 19 | VPCI 473/P/13 |  |  | India |  |
| 20 | 318043084 |  |  | Oman |  |
| 21 | 16B4865 | 16B15a |  | UK |  |
| 22 | VPCI 1133/P/13 | B11211 |  | India |  |
| 22 | VPCI 265/P/14 |  |  | India |  |
| 22 | VPCI 270/P/14 |  |  | India |  |
| 22 | VPCI 459/P/14 |  |  | India |  |
| 22 | VPCI 462/P/14 |  |  | India |  |
| 22 | VPCI 467/P/14 |  |  | India |  |
| 22 | VPCI 478/P/14 |  |  | India |  |
| 22 | VPCI 509/P/14 |  |  | India |  |
| 22 | VPCI 510/P/14 |  |  | India |  |
| 22 | 3420/15 |  |  | Kuwait |  |
| 22 | 4028/15 |  |  | Kuwait |  |
| 22 | 769/16 |  |  | Kuwait |  |
| 23 | VPCI 1132/P/13 | B11210 |  | India |  |
| 23 | VPCI 1134/P/13 |  |  | India |  |
| 23 | VPCI 250/P/14 | B11217 |  | India |  |
| 23 | VPCI 253/P/14 |  |  | India |  |
| 23 | VPCI 511/P/14 |  |  | India |  |
| 23 | VPCI 512/P/14 | B11214 |  | India |  |
| 23 | VPCI 513/P/14 | B11215 |  | India |  |
| 23 | VPCI 514/P/14 | B11216 |  | India |  |
| 23 | VPCI 550/P/14 |  |  | India |  |
| 23 | VPCI 674/P/12 |  |  | India |  |
| 23 | RCPF 1821 |  |  | Russia | Moscow |
| 23 | 15B34347 | 15B6 |  | UK |  |
| 23 | 15B42714 | 15B10 |  | UK |  |
| 23 | 15B54797 | 15B5 |  | UK |  |
| 23 | 16B10136 | 16B27b |  | UK |  |
| 23 | 16B1104 | 16B13 |  | UK |  |
| 23 | 16B264 | 16B12 |  | UK |  |
| 23 | 16B53197 |  |  | UK |  |
| 23 | 16B6373 | 16B20 |  | UK |  |
| 23 | 16B6920 | 16B21 |  | UK |  |
| 23 | 16B9044 | 16B24a |  | UK |  |
| 23 | 16B9260 | 16B22b |  | UK |  |
| 23 | 16B9331 | 16B15b |  | UK |  |
| 23 | 16B9828 | 16B25 |  | UK |  |
| 23 | 16B9856 | 16B26 |  | UK |  |
| 23 | 16I010622 |  |  | UK |  |
| 23 | 16I10156 |  |  | UK |  |
| 23 | 16I10532 | 16I30 |  | UK |  |
| 23 | 16I2986 | 16I29b |  | UK |  |
| 23 | 16I9766 |  |  | UK |  |
| 24 | VPCI 213/P/15 |  |  | India |  |
| 24 | VPCI 481/P/13 |  |  | India |  |
| 24 | VPCI 482/P/13 | B11207 |  | India |  |
| 24 | VPCI 67/P/15a |  |  | India |  |
| 24 | VPCI 714/P/14 |  |  | India |  |
| 24 | VPCI 73/P/15 |  |  | India |  |
| 25 | VPCI 900/P/15 |  |  | India |  |
| 26 | VPCI 1012/P/17 |  |  | India |  |
| 27 | VPCI 974/P/17-2 |  |  | India |  |
| 28 | CBS 15366 |  |  | Austria |  |
| 29 | 317052126 |  |  | Oman |  |
| 30 | CJ113 |  |  | Spain |  |
| 31 | CJ088 |  |  | Spain |  |
| 31 | CJ095 |  |  | Spain |  |
| 31 | CJ097 |  |  | Spain |  |
| 31 | CJ105 |  |  | Spain |  |
| 31 | CJ106 |  |  | Spain |  |
| 31 | CJ110 |  |  | Spain |  |
| 31 | CJ114 |  |  | Spain |  |
| 31 | CJ116 |  |  | Spain |  |
| 31 | CJ117 |  |  | Spain |  |
| 31 | CJ118 |  |  | Spain |  |
| 31 | CJ119 |  |  | Spain |  |
| 31 | CJ121 |  |  | Spain |  |
| 31 | CJ126 |  |  | Spain |  |
| 31 | CJ127 |  |  | Spain |  |
| 31 | CJ128 |  |  | Spain |  |
| 31 | CJ129 |  |  | Spain |  |
| 31 | CJ131 |  |  | Spain |  |
| 31 | CJ132 |  |  | Spain |  |
| 31 | CJ133 |  |  | Spain |  |
| 31 | CJ134 |  |  | Spain |  |
| 31 | CJ136 |  |  | Spain |  |
| 31 | CJ137 |  |  | Spain |  |
| 31 | CR180 |  |  | Spain |  |
| 31 | CR201 |  |  | Spain |  |
| 31 | CR312 |  |  | Spain |  |
| 31 | CR424 |  |  | Spain |  |
| 31 | CR531 |  |  | Spain |  |
| 32 | CJ124 |  |  | Spain |  |
| 33 | CJ130 |  |  | Spain |  |
| 34 | CJ096 |  |  | Spain |  |
| 34 | CJ098 |  |  | Spain |  |
| 34 | CJ115 |  |  | Spain |  |
| 35 | MOL208 | B11221 | CDC 383 | SA |  |
| 35 | MOL209-2 | B11222 | CDC 384 | SA |  |
| 35 | MOL224 | B11223 |  | SA |  |
| 35 | MOL293 | B11224 |  | SA |  |
| 35 | CJ094 |  |  | Spain |  |
| 35 | CJ099 |  |  | Spain |  |
| 35 | CJ100 |  |  | Spain |  |
| 35 | CJ102 |  |  | Spain |  |
| 35 | CJ103 |  |  | Spain |  |
| 35 | CJ104 |  |  | Spain |  |
| 35 | CJ107 |  |  | Spain |  |
| 35 | CJ109 |  |  | Spain |  |
| 35 | CJ109 |  |  | Spain |  |
| 35 | CJ111 |  |  | Spain |  |
| 35 | CJ112 |  |  | Spain |  |
| 35 | CJ120 |  |  | Spain |  |
| 35 | CJ140 |  |  | Spain |  |
| 35 | CR176 |  |  | Spain |  |
| 35 | CR184 |  |  | Spain |  |
| 35 | CR219 |  |  | Spain |  |
| 35 | CR220 |  |  | Spain |  |
| 35 | CR243 |  |  | Spain |  |
| 35 | CR244 |  |  | Spain |  |
| 35 | CR250 |  |  | Spain |  |
| 35 | CR309 |  |  | Spain |  |
| 36 | CR349 |  |  | Spain |  |
| 37 | CJ138 |  |  | Spain |  |
| 37 | CR157 |  |  | Spain |  |
| 37 | CR440 |  |  | Spain |  |
| 38 | KCTC 17810 |  |  | Korea |  |
| 39 | KCTC 17809 |  |  | Korea |  |
| 40 | JCM 15448 | B11220 | DSMZ 21092 | Japan |  |
